# Evaluating spatiotemporal integration of shape cues

**DOI:** 10.1101/809103

**Authors:** Taylor Burchfield, Ernest Greene

**Affiliations:** Laboratory for Neurometric Research, Department of Psychology, University of Southern California, Los Angeles, California 90089 USA

## Abstract

Prior work has shown that humans can successfully identify letters that are constructed with a sparse array of dots, wherein the dot pattern reflects the strokes that would normally be used to fashion a given letter. In the present work the dots were briefly displayed, one at a time in sequence, varying the spatial order in which they were shown. A forward sequence was spatially ordered as though one were passing a stroke across the dots to connect them. Experiments compared this baseline condition to the following three conditions: a) the dot sequence was spatially ordered, but in the reverse direction from how letter strokes might normally be written; b) the dots in each stroke of the letter were displayed in a random order; c) the sequence of displayed dots were chosen for display from any location in the letter. Significant differences were found between the baseline condition and all three of the comparison conditions, with letter recognition being far worse for the random conditions than for conditions that provided consistent spatial ordering of dot sequences. These findings show that spatial order is critical for integration of shape cues that have been sequentially displayed.

## INTRODUCTION

Experience dictates that humans can more readily perceive objects when the boundary cues are displayed an orderly, systematic manner. It is likely that natural camouflage is effective because information about the object’s boundaries are spatially ambiguous and intermittent —as can occur when an animal moves behind vegetation. The mechanisms providing for shape recognition often draw on the concept of closure proposed by Gestalt psychologists, wherein the parts of an object contribute to generating an integrated whole. When this integration is disrupted by inconsistent presentation of the shape cues, one is less likely to recognize that object (Elder, 2018).

Prior work has demonstrated that the human visual system is capable of perceiving shapes with minimal stimulus information, such as when the shape is represented by a pattern of dots and the display provides only a sparse “low-density” sample of the pattern (Greene & Hautus, 2018; Nordberg, Hautus, & Greene, 2018; Greene & Visani, 2015). For letters formed as single-file strings of dots, respondents manifest above chance recognition of the letters at 3% density. The performance is near-perfect at 27% density, i.e., wherein roughly every fifth dot in the string is included in the letter display (Greene, 2016a). This suggests that the information provided by the strokes normally used to create letters is highly redundant. The discrete strings of dots from which a low-density sample is drawn already has gaps between each dot, yet one can eliminate four out of five dots and still see near perfect recognition of the letters.

Humans have also been shown to be able to integrate shape information across an interval of at least 200 ms. This can be attributed to the phenomenon of information persistence (Coltheart, 1980; Greene & Visani, 2015). The persistence allows the visual system to retain stimulus information for a sizeable temporal interval, allowing it to be integrated with related cues that are subsequently provided.

Given that the visual system can identify shapes from minimal cues and with temporal separation of those cues, the goal of this work is to more fully investigate how the sequencing of the cues affects the integration process. One might reasonably assume that systematic presentation of boundaries information would be essential to integration. Shipley and Kellman (1996) examined the role of ordered display of shape elements with a deletion/accretion paradigm, wherein a shape moves behind an occluding surface, sequentially modifying visibility of boundary cues. They confirmed that the human perceptual system contains mechanisms that compensate for fragmented boundary presentation. A subsequent study by McCarthy, Erlikhman, and Caplovitz (2017) examined how the perception of an ambiguous stimulus can be illuminated within a moving zone that frames the stimulus, i.e. a “spotlight.” The fluid illumination of different aspects of a scene allows the observer to determine what shapes are present.

Here we use sequential displays of low dot-density letters to investigate the integration of stimulus information as a function of spatial positioning and temporal separation. Letters were used because they can be reliably created and identified from successive graphic strokes. Properly speaking, a “stroke” refers to a letter element that is continuous while being written. For example, an A has three strokes, two that are diagonal and one that connects the diagonals near the center; a B consists of a vertical stroke and two curved strokes. The spaced dots of our displays were positioned such that a continuous stroke could be drawn to connect them. It is convenient to describe the spatial relationships as strokes, even though the dots were not connected. [See Methods for details on the operational definition of stroke-sequences used in the present work.]

The left panels of Figure 1 illustrates the strings of adjacent dots that would comprise a given letter at 100% density, and the right panels show examples at 30% density. In each of the three experiments described below, a baseline condition was sequential display of dots that followed the path that would produce a handwritten letter. The dots were briefly displayed one at a time, as though an unseen stroke was causing each to be briefly illuminated. Further, the order of successive strokes themselves matched how the letter could be graphically generated [See Methods.] This baseline condition can also be described as the “forward” treatment condition, meaning that the display sequence followed the normal direction for writing each of the several successive strokes for a given letter.

**Fig 1.**
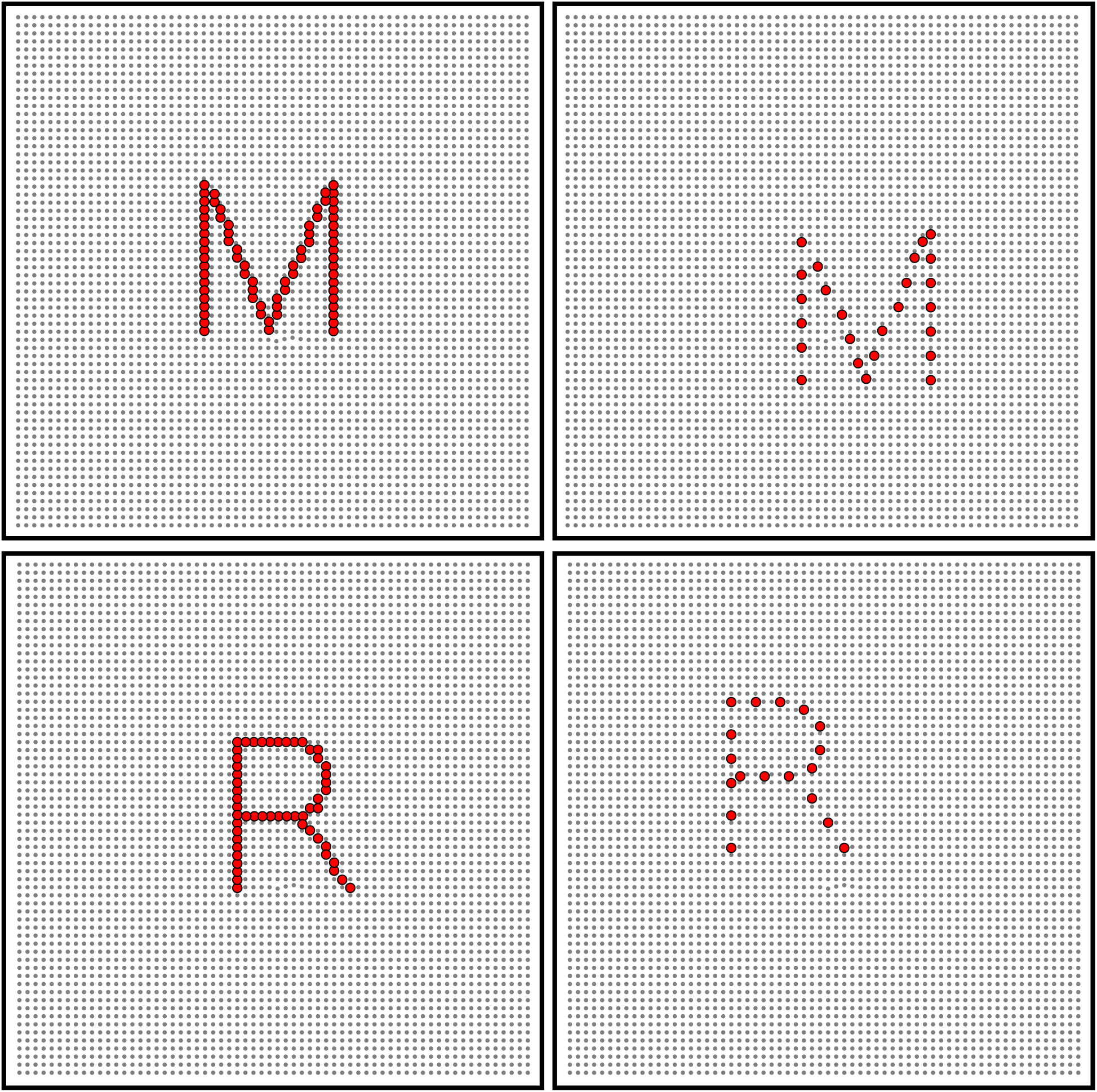
Illustration of dot density condition. The left panels show examples wherein the array of dots forming the letter strokes are at 100% density. The right panels show the low-density (30%) versions, wherein the dots were briefly displayed one at a time, with the order of display being varied.

Experiment 1 compared the baseline condition with a “reverse” condition, i.e., the dots were displayed in the opposite order with respect to the baseline condition. This reversed not only the order within a given stroke sequence, but the strokes themselves. Experiment 2 compared the baseline condition with a “random within strokes” condition. Here the order of strokes was the same as baseline, but the order of dot display within strokes was random. Experiment 3 compared the baseline condition with a “completely random” condition, where the order of successive dots was chosen at random from the low-density letter, irrespective of stroke membership.

## RESULTS

Each of the experimental treatments was designed to produce a decline in recognition as a function of the amount of delay between successive dot displays. Mixed effect logistic regression confirmed a significant decline for each treatment at p < 0.0001 in each of the experiments. The linear, quadratic, and cubic components that were significant for a given treatment provide models of treatment effects in the following figures, along with 95% confidence bands.

Experiment 1 evaluated how the interval between successive dots would affect letter recognition, comparing the effects of a forward sequence in relation to a backward sequence. The resulting models both show a dominant linear decline in recognition, with the rate of change being roughly the same for both treatments (see Figure 2). Evaluating the treatment levels individually, the impairment was significant at 50 ms of dot separation and longer. The consistent differential in probability of recognition across all intervals greater than zero suggests that a forward sequence is better able to elicit memory of letter attributes. This may be related to the normal direction of eye scans in reading of written English, as will be discussed subsequently.

**Fig 2.**
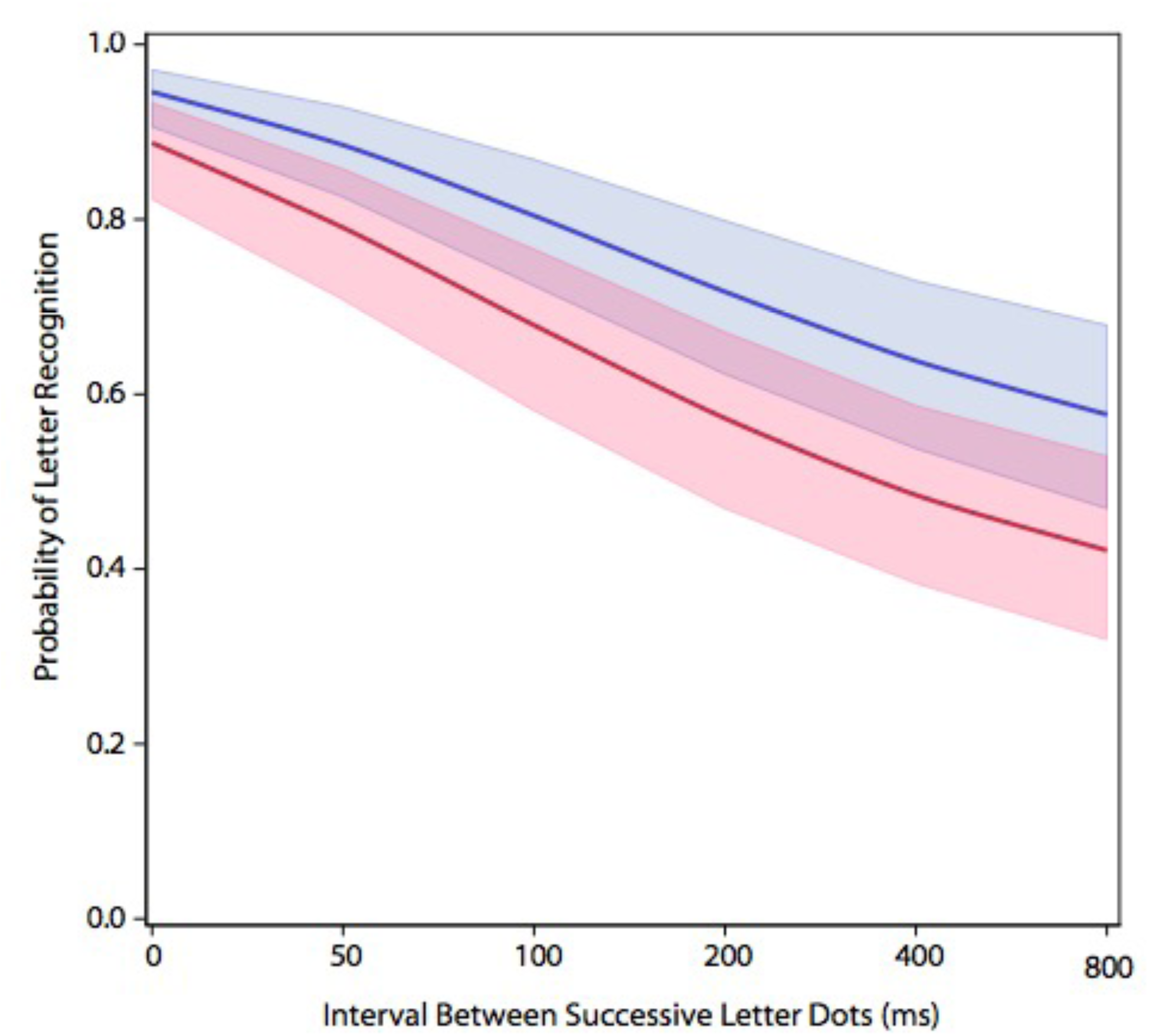
Comparing a forward dot sequence to a backward dot sequence. Presenting the letter dots in a backward sequence (red) produced a larger decline in letter recognition than was produced by the forward sequence (blue) – p < 0.0001.

Experiment 2 evaluated letter recognition where the sequential dots were chosen at random from the full letter pattern, compared to presentation of dots using a forward sequence. One can see that the resulting model for the random condition adds a very strong quadratic component to the model, with recognition sinking to about 40% recognition with only 50 ms of dot separation (see Figure 3). Overall, the impairment of recognition for the random sequence can be seen to be substantially larger than was found in Experiment 1. This differential is further analyzed below.

**Fig 3.**
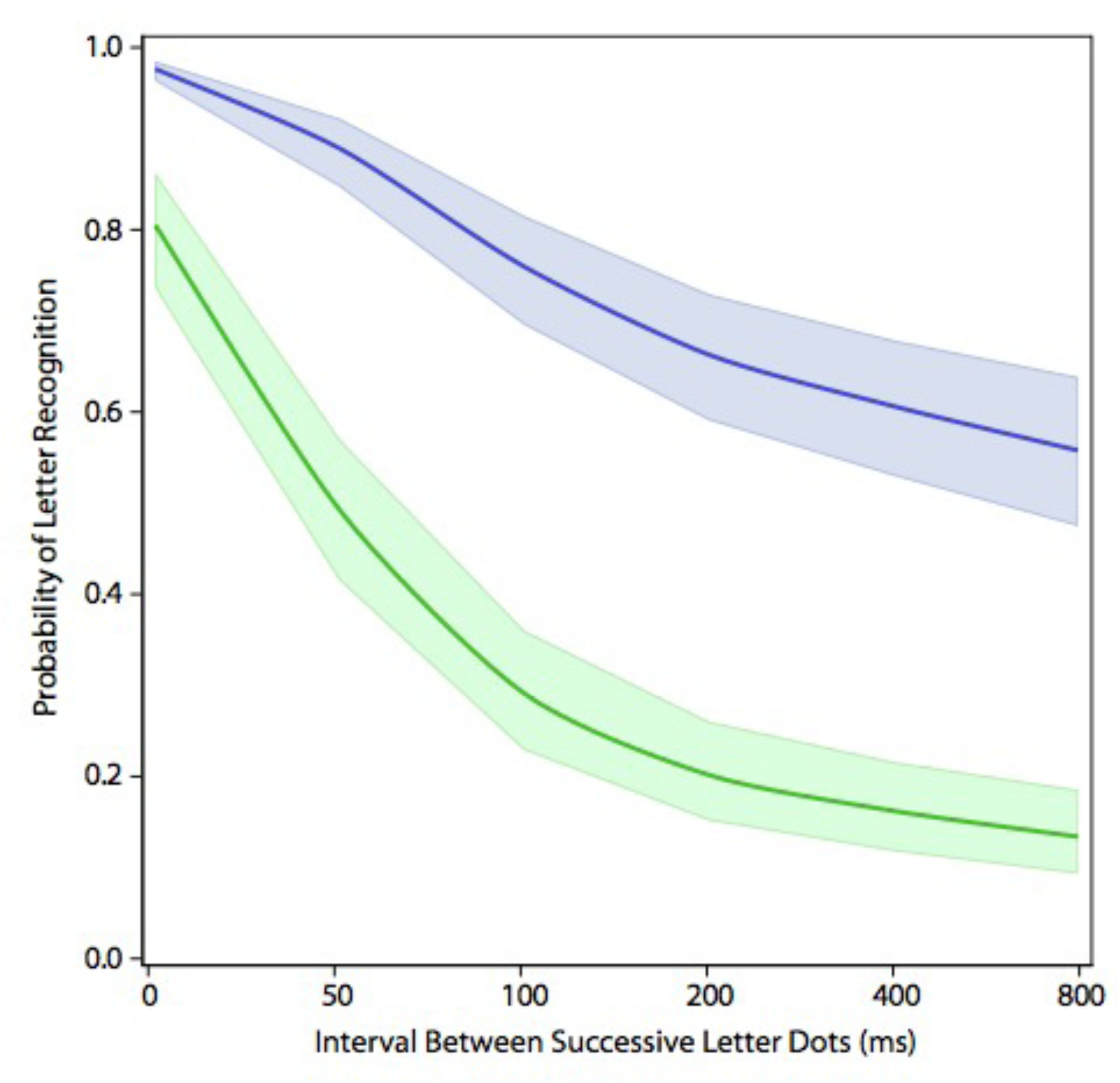
Comparing a random dot sequence to forward dot sequence. Presenting the letter dots in a random sequence (green) produced a larger decline in letter recognition than was produced by the forward sequence (blue) – p < 0.0001.

Experiment 3 compared the forward condition against a condition wherein the letter strokes were presented in the same order as the forward treatment condition, but with dots within each stroke being chosen in a random order. One respondent had mean letter recognition for the simultaneous (SOA = 0) display condition that was p < 0.0001 below the group mean, so the statistical analysis of treatment effect was done without including data from this respondent. The resulting model for the “random-within-stroke” condition also had large non-linear components, as can be seen in Figure 4. Here also, the overall amount of impairment was well below that produced by forward sequencing of dots.

**Fig 4.**
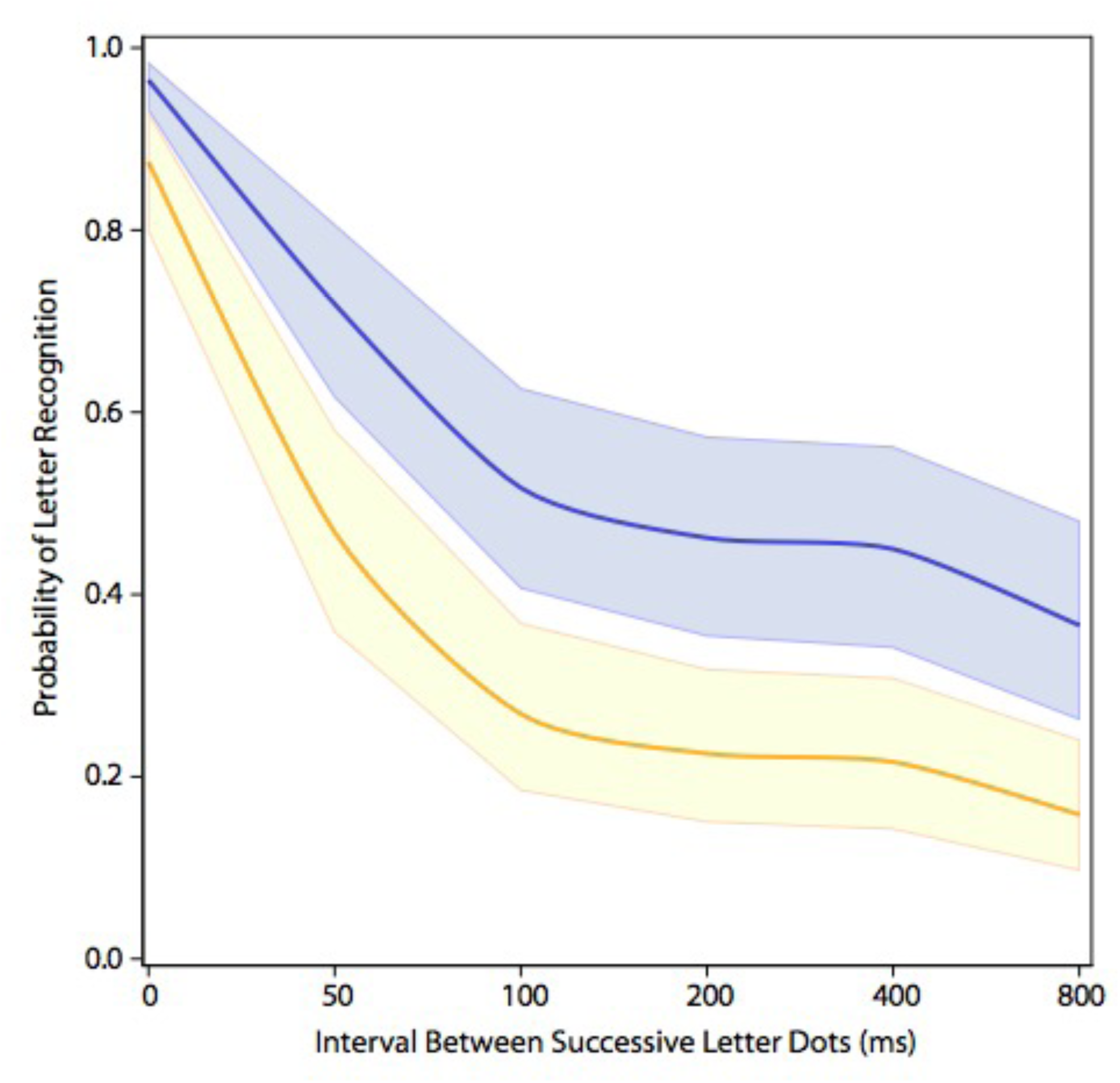
Comparing a random within-stroke dot sequence to a forward sequence. Presenting the letter dots in a random sequence within each stroke (gold) produced a larger decline in letter recognition than was produced by the forward sequence (blue) – p < 0.0001.

Presentation of dots as a forward sequence was the baseline treatment for each of the three experiments. The relative influence of non-baseline conditions was of special interest. The backward sequence (Exp 1, red) differed significantly from the totally random sequence (Exp 2, green) and from the random-within-strokes condition (Exp 3, gold), with p < 0.0001 for each comparison. However, the totally random sequence (Exp 2, green) did not significantly differ from the random-within-strokes condition (Exp 3, gold), with p < 0.06289. Models showing the amount of recognition impairment for the latter comparison are illustrated in Figure 5. It seems clear that sequential display of dots selected from random locations greatly impairs letter recognition, and does so even if one has limited the choice to dots that all lie within a given stroke. The strokes that comprise a given letter are essentially its contours, and these results may well pertain to the broader topic of how the contours of diverse objects are registered for purposes of object recognition.

**Fig 5.**
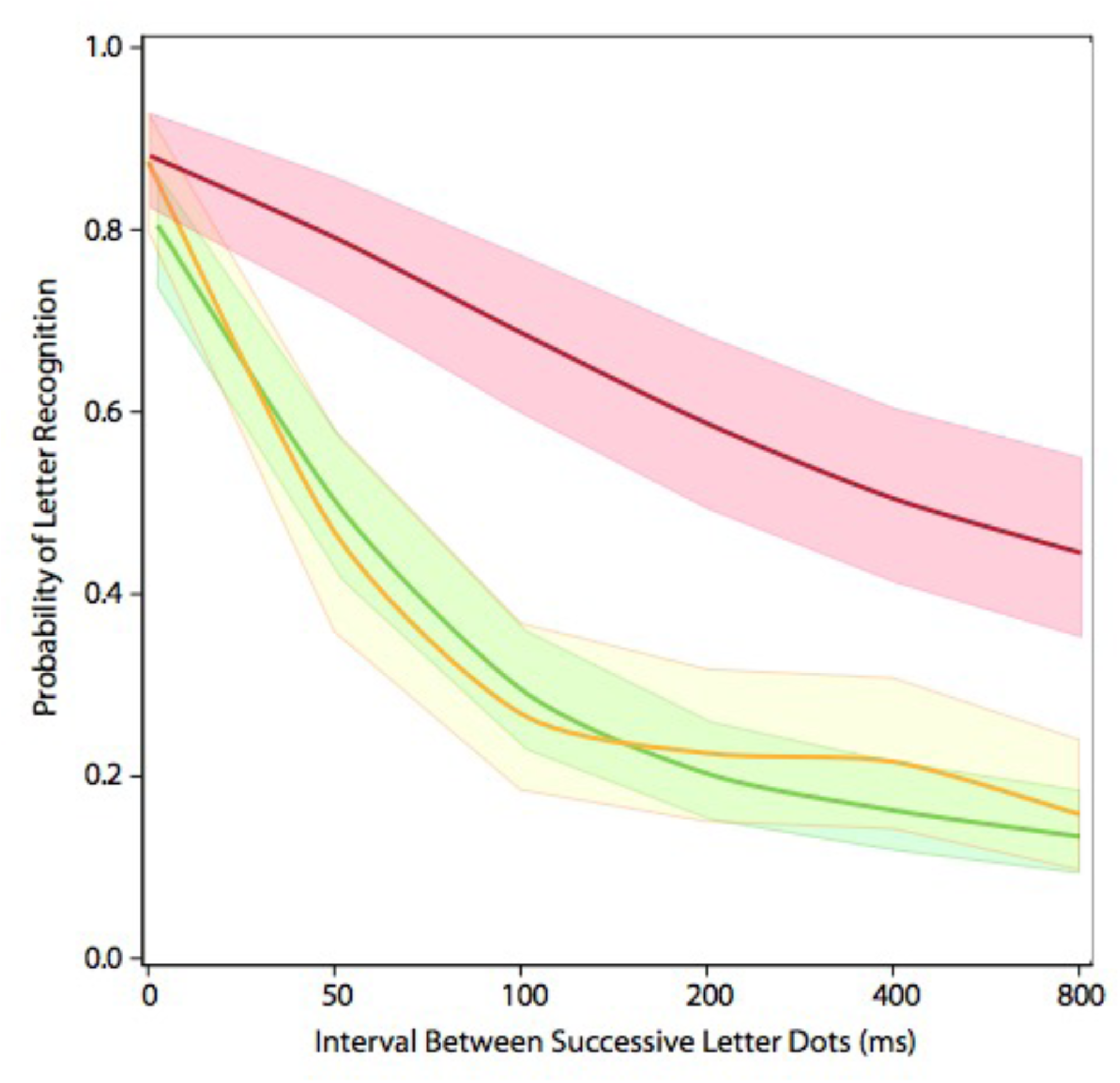
Comparing non-ordered sequences to the backward sequence. The presentation of successive dots as a backward sequence (Exp 1, red) differed significantly for the other two non-ordered conditions – p < 0.0001 for each comparison. The random-within-strokes condition (Exp 3, gold) did not differ significantly from the totally random sequencing of dot display (Exp 2, green).

To be thorough, the forward sequence treatments for the three experiments (blue in Figures 1-3) were statistically examined. That condition did not produce a significant differential in recognition for the groups tested in Experiment 1 versus Experiment 2 (p = 0.8710). However, the forward model in Experiment 3 was significantly different from Experiment 1 (p < 0180) and from Experiment 2 (p < 0.0290). We have no explanation for why subjects of Experiment 3 had lower letter recognition when provided with the same dot sequence used in the first two experiments.

## DISCUSSION

The three experiments demonstrated a significant difference between the baseline (forward) order of dot presentation and the other display orders that were evaluated. We therefore suggest that the forward sequencing of dots can be integrated more readily, allowing registration of the strokes on which the memory of letter-shape is based. This could be due to a lifetime of experience with writing letters as a forward sequence of strokes.

Further, we found that backward strokes yielded better recognition than either of the random conditions, which reflects an advantage for spatial ordering of the stroke markers. The ability to integrate the cues across time requires persistence of shape information, and the results show that visual system can better integrate a temporal sequence if the dots are adjacent. In other words, as successive dost are briefly flashed, their locations persist in the visual system for a short period of time. Display of adjacent locations within the dot pattern contributes to registering the stroke, and thus the letter. So when the dots are presented in a consistent spatial order, the observer is able to better synthesize the stimuli, and can therefore perceive a more cohesive stimulus.

Visible persistence, which lies at the retinal level, can last for about a hundred milliseconds, whereas cortically based information persistence can provide for above-chance recognition for up to seven hundred milliseconds (Greene & Visani, 2015; Greene 2016a). With these time periods in mind, we can speculate about the role of persistence in the integration of the present display sequences. At 30% density, the mean number of dots per stroke is just over six dots. A fifty millisecond interval between successive dot would allow fourteen dots to last in working memory and a hundred millisecond interval would allow seven dots to be perceived. Therefore, most or all of the dots in a given stroke would be available to working memory. This could account for the minimal differential between forward and backward treatment conditions. However, it presents a challenge to the dramatically impaired performance for the random-within-stroke and random-within-shape conditions that were manifested with a temporal separation of only fifty milliseconds.

Information persistence failed to provide for recognition not only when the random selection of dots was from all portions of the letter, but also where the random choice was within a given stroke. The recognition deficits produced by the random display conditions can be attributed to “ineffective spatiotemporal integration,” in this case a failure to integrate brief displays of non-adjacent spatial information. These conditions prevent recognition of the letter shape even though all of the stimulus cues are simultaneously available to working memory. It is as though random presentation of the dots “cancels out” preceding and succeeding shape cues, which precludes synthesis of the cues, and thus recognition.

Treisman (1986) described this idea when investigating the phenomenon of illusory conjunctions. She found that objects having the same color, shape, or proximity to other similar objects yielded an illusory distinction between those items. As applied to the present displays, dots that were randomly ordered might be incorrectly paired, leading the observer to commit an error in processing the configuration. She related this phenomenon to the Gestalt principles of grouping, wherein the sum of parts would be expected to form a cohesive whole.

Another relevant process that could apply here is the accretion/deletion of shape cues. In nature, animate (moving) objects are often occluded by other elements of the scene, producing systematic, predictable sequences of boundary cues. Presenting one dot at a time in sequence is similar to boundary cues being “deleted;” as an object moves behind an occluder. Shipley and Kellman (1997) found that when objects moved through a complex scene with fragmented boundaries, recognition of the object declined as a function of the degree of fragmentation. In this study, they increased the number of target fragments, but not the number of noise fragments. They found that when there is a higher ratio of noise fragments relative to target fragments, recognition declined. They concluded that the extraneous elements from the scene were being incorporated into motion tracking mechanisms, which disrupted the accretion-deletion process. From this perspective, the ineffective spatiotemporal integration that our experimental conditions produced could be seen as an impairment of accretion/deletion mechanisms.

Related to accretion/deletion is perception of a scene through a moving slit. The ordered stroke condition most closely mimics the presentation of dots as seen through a moving slit—as the majority of the letter is concealed, the slit allows the observer to uncover shape elements in a systematic manner. Rock and Sigman (1973) found that when a moving slit was presented over a line figure, participants are able to perceive a cohesive shape. Briefly presenting the stroke dots in a consistent spatial order emulates the temporary nature of seeing the shape boundary through a moving slit.

Even though the random-within-strokes condition presented strokes individually, it did not present the letters in a way that was “traceable;” i.e. it did not emulate how an object would appear and reappear in the environment, for it lacked continuity. Scholl, Pylyshyn, and Feldman (2001) found that object tracking is greatly enhanced when subjects can easily “pair” stimuli, i.e. when they take the same trajectory across a scene. They also found that ambiguous ends of stimuli, such as those that seemed to change in length, were not treated as objects because they were harder to pinpoint and track. Interestingly, they discovered that tracking was impeded by non-moving objects that were present in the background. Thus, it seems as if the mere number of objects in the area of focus has an impact on stimulus tracking. It is certainly possible that the accretion/deletion phenomenon presents the shape in a way that is conducive to the appearance of a single object (a stroke), and is thus easier to recognize. One continuous sequence of adjacent dots is more like a single object than a random presentation of dots, and can be perceived as being occluded.

Overall, the present results support the view that the visual system can encode sequential shape cues more effectively if they are spatially ordered. This is consistent with prior proposals that the initial stages of shape encoding are accomplished in the retina and/or superior colliculus, using scan waves to register the relative location of boundary markers (Greene, 2007, 2016b, 2018; Greene & Morrison, 2018; Greene & Patel, 2018). This concept views elemental encoding of shapes as having evolved from primitive mechanisms for motion analysis, wherein successive portions of the shape’s boundary are registered as they pass across the retina. The process converts a two-dimensional shape into a one-dimensional signal that can be stored and used for subsequent recognition. Further evolution of this mechanism would yield the ability to generate scan waves that could produce the same kind of summary message in the presence of still images, i.e., where the shape to be recognized did not move. According to this view, effective recognition of a shape would require delivery of a spatially ordered sequence of shape cues, as was found here.

## METHODS

### Authorization, Consent, and Participation

The protocols for these experiments were approved by the USC Institutional Review Board. Respondents were recruited from the Psychology Subject Pool. Each respondent provided informed consent for being tested, which included provisions for termination of testing without penalty upon request by the respondent. A total of 24 respondents provided the data, eight in each of the three experiments.

### Display Equipment

Letters were displayed as brief sequential flashes from a 64×64 array of LEDs (dots) mounted on a display board. Respondents viewed the display board at a distance of 3.5 m, and at this distance the visual angle of a given dot was 4.92 arc’, dot to dot spacing was 9.23 arc’, and the total span of the array (edge to edge) was 9.80 arc°. Ambient illumination of the room was 10 lux.

### Letter Configurations

A table of dot locations represented letters as strings of adjacent dots, each letter being 20 dots tall and with a maximum width of 14 dots. It is convenient to describe the dot sequences as “strokes,” with the stroke sequences being configured as they might be written by hand. These stroke sequences are illustrated in Figure 6. For example, the letter A was specified as having three strokes, starting at the apex and proceeding down the left stroke, returning to the apex and proceeding down the right stroke, then starting at the middle of the left stroke and passing to the right to create the horizontal stroke. The letter B would be specified as beginning at the top dot to create the vertical stroke, then returning to the top to produce the upper loop-stroke, followed by the lower loop-stroke. The C was considered to be a single stroke, starting at the top-end dot and sequencing through each connected dot to end at the bottom end. Figure 6 illustrates the stroke specification for each letter of the alphabet.

**Fig 6.**
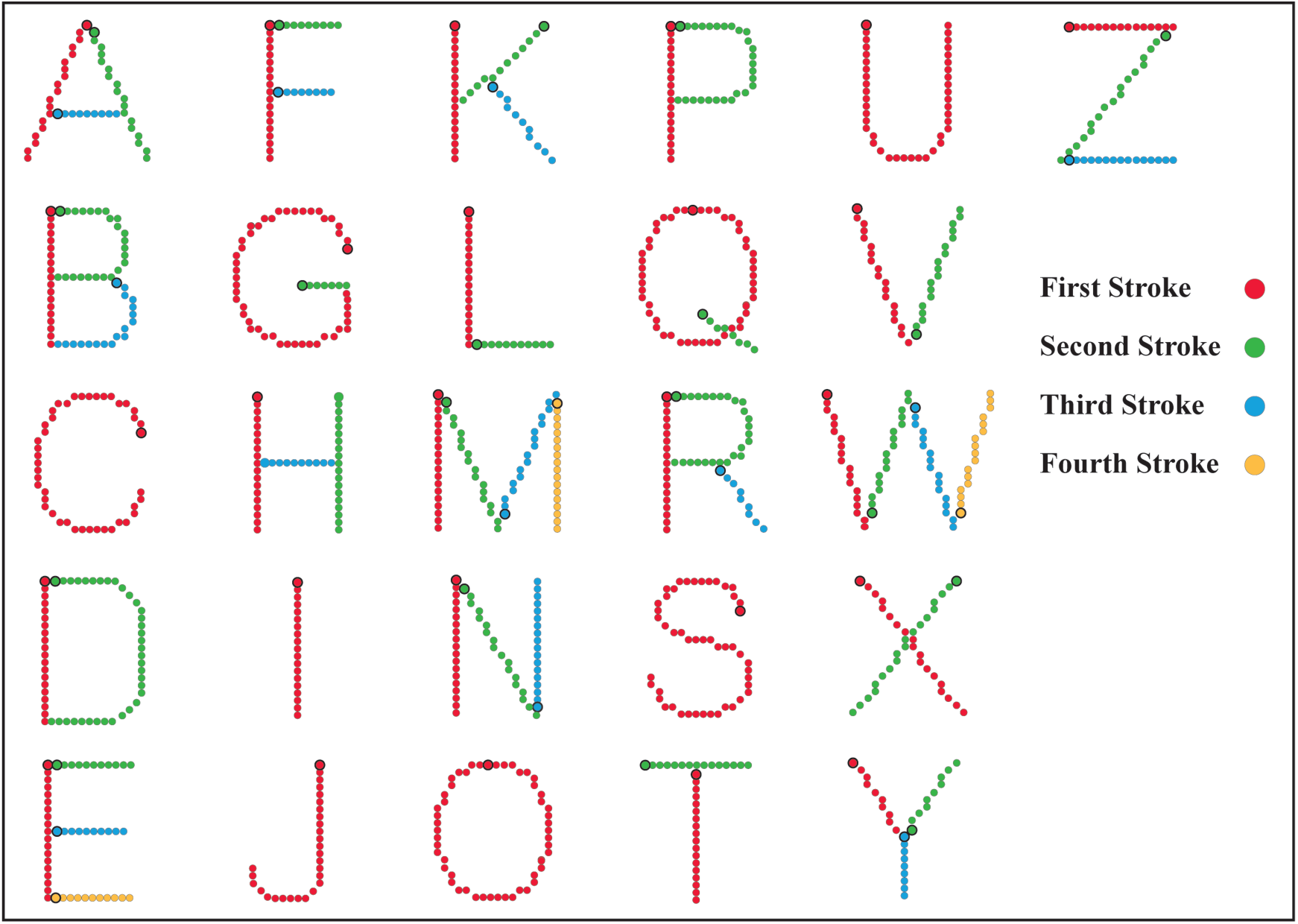
Alphabet showing the forward order of stroke sequences. The table of addresses represented letters of the alphabet as strings of adjacent dots (strokes), with most letters being constructed with several strokes. This illustration shows all the dots there were available in the address table for the letters, but the displays themselves provided a low-density sampling of the dots. For the baseline (forward) condition used in each of the three experiments, dots were presented in the order specified by the color code of this illustration. For a given stroke, the sequence began at the dot with a dark boundary. Using the letter Z as the example, the first stroke sequence (red) begins with the left dot; the second stroke sequence (green) begins at the top of the diagonal; the third stroke sequence (blue) begins with the left dot.

In each of the three experiments, the letter pattern that was displayed on a given trial was a reduced-density (30%) sample of the full complement of dots available in the letter. Prior research (Greene, 2016) had found recognition to be asymptotic, near 100%, at higher dot densities. A 30% sample was expected to provide for 80-90% recognition when all the dots were displayed at the same moment, and the goal was to provide a decline from that high level as the time interval between successive dots was increased (see timing conditions, below). For each letter displayed, the 30% sample was chosen at random, on the fly, but with an algorithm designed to maximize spacing of adjacent dots within a given stroke. It is convenient to describe the 30% sample as a “stroke” even though it provides only some of the dots contained within the table of address locations.

### Letter Display Conditions

On each trial in each of the three experiments, positioning of a given letter on the display board was varied at random, with horizontal and vertical offsets of up to five dots. For a given display, the dots of the 30% sample were sequentially displayed, each as an ultra-brief flash at an intensity of 1000 µW/sr for a duration of 10 µs. The stimulus onset interval (SOA) between successive dots was varied across six levels, these being: 0, 50, 100, 200, 400, and 800 milliseconds. A randomly selected letter was displayed at each of these treatment levels 30 times, for a total of 360 total trials. The experimental condition that distinguished the experiments was the order in which dots were selected for display, as follows.

Experiment 1 compared the “forward” condition with a “backward” condition. The forward condition ordered the dot sequence as illustrated in Figure 6, this being a baseline condition for each of the three experiments. For the “backward” condition the address list for the letter was read in reverse order, which not only reversed the sequence within each stroke but also the order in which strokes were delivered.

Experiment 2 compared the “forward” condition to a “random” condition that displayed 30% of the dots from various dot locations within a given letter. The random selection of dot locations was done “on the fly” during testing, thus providing each subject with a different order of dot locations.

Experiment 3 compared the “forward” condition with a “random within stroke” condition, wherein the random choice was among the 30% sample of stroke dots, with the order of strokes being the same as for the forward condition.

### Experimental Test Conditions

After receiving consent instructions, the respondent was seated against a wall of the test room, facing the display board, which was mounted on the opposite wall at eye level. A single pulse-width modulated light bulb was mounted about one meter above the respondent’s head, this providing ambient light for the room.

Each trial was preceded by a fixation marker, i.e., four dots at the center of the board emitting light at an intensity of 0.2 µW/sr. This marker disappeared immediately before display of the letter dots. The respondent was expected to say what letter had been displayed, and to guess if they did not recognize it or were unsure. In general this response was provided within 1-2 seconds. The test administrator then entered the letter that was named through a keyboard, which logged the response, along with specifics about which letter had been displayed and the treatment conditions for that trial. Entry of this information immediately launched the next trial, beginning with display of the fixation marker for half a second, followed by display of the next letter. Presentation of all trials generally took 40-45 minutes, and all respondents who were recruited for testing were able to complete all display trials.

## Acknowledgments

The display equipment was designed and fabricated by, and experimental applications were written by Jack Morrison, Digital Insight. Dr. Wei Wang, Harvard Medical School provided statistical analysis and modeling of data. Funding for this research was provided by the Neuropsychology Foundation and the Quest for Truth Foundation.

## Notes

#### Summary of Updates

Replacing initial submission with a version that has figures included with the text.

## REFERENCES

Coltheart, M., Iconic memory and visible persistence. Percept Psychophys, (1980) 27, 183–228.

Elder, J. H., Shape from Contour: Computation and Representation. Annu Rev Vis Sci, 4(1), 423–450. (2018) doi:10.1146/annurev-vision-091517-034110

Froyen, V., Feldman, J., & Singh, M., Rotating columns: Relating structure-from-motion, accretion/deletion, and figure/ground. J Vis, 13(10), 6–6 (2013). doi:10.1167/13.10.6

Greene, E., Retinal encoding of ultrabrief shape recognition cues. PLOS ONE, 2(9), e871 (2007)

Greene, E., Information persistence evaluated with low-density dot patterns. Acta Psychol, 170, 215–225 (2016a). doi:10.1016/j.actpsy.2016.08.005

Greene, E., How do we know whether three dots form an equilateral triangle? JSM Brain Sci, 1(1), (2016b) 1002.

Greene, E., New encoding concepts for shape recognition are needed. AIMS Neurosci, 5 (2018), 162–178.

Greene, E., & Morrison, J., Computational scaling of shape similarity that has potential for neuromorphic implementation. IEEE Access, 6 (2018) 38294–38302.

Greene, E., & Hautus, M.J., Evaluating persistence of shape information using a matching protocol. AIMS Neurosci, 5(1) (2018), 81–96. doi:10.3934/neuroscience.2018.1.81

Greene, E., & Patel, Y., Scan transcription of two-dimensional shapes as an alternative neuromorphic concept. Trends Artif Intel, 1(1) (2018) 27–33.

Greene, E., & Visani, A., Recognition of letters displayed as briefly flashed dot patterns. Attn, Percept, Psychophys, 77(6) (2015), 1955–1969. doi:10.3758/s13414-015-0913-6

Mccarthy, J. D., Erlikhman, G., & Caplovitz, G. P., The maintenance and updating of representations of no longer visible objects and their parts. Progress Brain Res Temp Sampl Represen Updat, 163–192 (2017). doi:10.1016/bs.pbr.2017.07.010

Nordberg, H., Hautus, M. J., & Greene, E., Visual encoding of partial unknown shape boundaries. AIMS Neurosci, 5(2) (2018), 132–147. doi:10.3934/neuroscience.2018.2.132

Scholl, B. J., Pylyshyn, Z. W., & Feldman, J.,. What is a visual object? Evidence from target merging in multiple object tracking. Cognition, 80(1-2) (2001), 159–177. doi:10.1016/s0010-0277(00)00157-8

Shipley, T. F., & Kellman, P. J., Spatio-temporal boundary formation: The role of local motion signals in boundary perception. Vis Res, 37(10) (1997), 1281–1293. doi:10.1016/s0042-6989(96)00272-6

Rock, I & Sigman, E., Factors in the Perception of Form through a Moving Slit. Sage J, 2(3) (1973), doi: 10.1068/p020357.

Treisman, A., Features and Objects in Visual Processing. Sci Am, 255(5) (1986), 114–125.

